# NSF is required for diverse endocytic modes by promoting fusion and fission pore closure in secretory cells

**DOI:** 10.1101/2025.08.13.670133

**Authors:** Xin-Sheng Wu, Tao Sun, Bo Shi, Sunghoon Lee, Zheng Zhang, Lisi Wei, Xin Wang, Maryam Molakarimi, Sue Han, Aaron Uy, Lin Gan, Ling-Gang Wu

## Abstract

The ATPase N-ethylmaleimide-sensitive factor (NSF), known for disassembling SNARE complexes, plays key roles in neurotransmitter release, neurotransmitter (AMPA, GABA, dopamine) receptor trafficking, and synaptic plasticity, and its dysfunction or mutation is linked to neurological disorders. These roles are largely attributed to SNARE-mediated exocytosis. Here, we reveal a previously unrecognized role for NSF: mediating diverse modes of endocytosis—including slow, fast, ultrafast, overshoot, and bulk—by driving closure of both fusion and fission pores. This function was consistently observed across large calyx nerve terminals, small hippocampal boutons, and chromaffin cells using capacitance recordings, synaptopHluorin imaging, electron microscopy, and multi-color pore-closure imaging. Results were robust across four NSF inhibitors, gene knockout, knockdown, and specific mutations. These findings establish NSF as a central regulator of membrane fission, kiss-and-run fusion, endocytosis, and exo-endocytosis coupling—offering new mechanistic insights into its diverse physiological and pathological roles in synaptic transmission, receptor trafficking, and neurological diseases.

## Introduction

The ATPase N-ethylmaleimide-sensitive factor (NSF) is well-known to disassemble the soluble NSF attachment protein receptor (SNARE) complex, composed of synaptobrevin, SNAP-25, and syntaxin, which drives vesicle fusion, releasing neurotransmitters and hormones. Genetic loss of NSF results in the accumulation of *cis*-SNARE complexes on synaptic vesicles and inhibition of transmitter release in Drosophila synapses^1^. Acute peptide-based perturbation experiments in giant squid synapses show that impaired NSF activity causes a rapid block in neurotransmission and slowing down of release, suggesting that SNARE complex disassembly is crucial for transmitter release at the pre-fusion state^2,3^. Mutation and pharmacological inhibition of NSF reveal NSF in regulating vesicle priming for fusion at release sites in Drosophila synapses and mouse hippocampal synapses^4,5^, suggesting that the SNARE complex disassembly is important for vesicle priming at the pre-fusion state. Besides regulating vesicle fusion at the nerve terminal, NSF has been shown to regulate the trafficking of important neurotransmitter receptors, such as AMPA receptors, GABA receptors, and dopamine receptors, thereby regulating synaptic plasticity. Aggregation, functional impairment, and/or mutation of NSF have been linked to major neurological disorders, including Parkinson’s disease, Alzheimer’s disease, and epilepsy^6^. In brief, NSF is crucial for transmitter release, neurotransmitter receptor trafficking, synaptic plasticity, and neurological disorders. These diverse physiological functions and pathological roles have been largely attributed to its ability to disassemble the SNARE complex, crucial for vesicle priming and release^6^.

Little is known about whether NSF plays any role after vesicle fusion, such as the fusion pore expansion and closure, and subsequent endocytosis that recycles exocytosed vesicles to sustain synaptic transmission and exocytosis in secretory cells. Here, we studied whether NSF is involved in these post-fusion roles by 1) measuring slow, fast, and ultrafast endocytosis with capacitance measurements in chromaffin cells and calyx of Held synapses, 2) detecting endocytosis with synaptopHluorin imaging in hippocampal synapses, and 3) quantifying bulk endocytosis with electron microscopy in hippocampal synapses. By inhibiting NSF with various pharmacological inhibitors, gene knockout or knockdown, we found, to our surprise, that NSF is essential for each of these different forms of endocytosis in three preparations. By imaging fusion pore opening and closure, as well as the fission pore closure, we found that NSF is crucial in mediating diverse forms of endocytosis mentioned above by playing a critical role in closing both fusion and fission pores in chromaffin cells. These results suggest including NSF as a key player in the current models of membrane fission, diverse modes of endocytosis, vesicle recycling, and exo-endocytosis coupling, which may contribute to accounting for NSF’s diverse physiological and pathological roles discussed above.

## Results

### NSF is involved in slow and fast endocytosis at calyces

#### Recording conditions

We measured slow (Fig. 1A-E) and fast (Fig. 1F-J) endocytosis by whole-cell capacitance recordings at calyces with a pipette containing either a control solution or one of the following four types of NSF (ATPase) inhibitors: 1) ATPγS (replacing ATP, 4 mM, n = 6) or 0 ATP (n = 7), 2) N-ethylmaleimide (NEM, 1 mM, n = 12), 3) an NSF peptide (NSF_p_, 1 mM, n = 9)^3^, and 4) a SNAP peptide (SNAP_p_, 1 mM, n = 8) that blocks the binding between NSF and SNAP (soluble NSF attachment protein), a chaperon for recruiting NSF^7^. The corresponding control solution for peptides contained either mutated NSF_p_ (NSF_mp_, 1 mM, n = 10) or scrambled SNAP_p_ (SNAP_sp_, 1 mM, n = 7).

**Figure 1.**
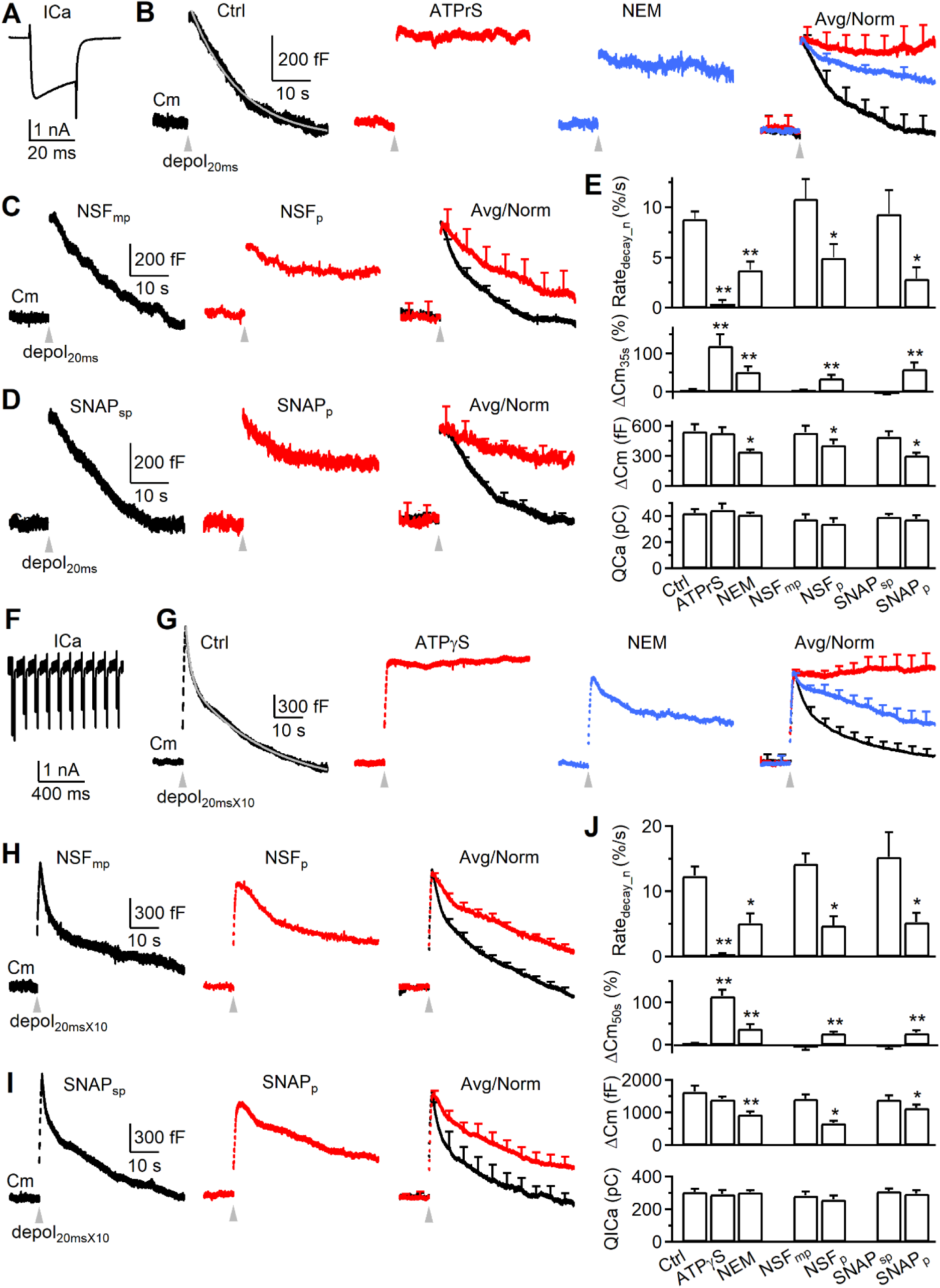
NSF is involved in slow endocytosis at calyces. (A) Sampled calcium current (ICa) induced by depol_20ms_ in a calyx of Held. (B) Sampled (single traces, left three traces) or averaged (right) capacitance changes induced by depol_20ms_ (grey arrow head) at 4 - 10 min after break-in with a pipette containing a control solution (black, n = 11), 4 mM ATPγS (replacing ATP, red, n = 6) or 1 mM NEM (blue, n = 12). The Cm decay in Ctrl was fit mono-exponentially (τ: 10.6 s, gray). The amplitude of the ΔCm_peak_ for averaged traces was normalized (Avg/norm), and data were expressed as mean + SE every 5 - 10 s (applies to all other panels). Scale bars apply to panels B-D. (C) Similar to panel B, but with NSF_mp_ (1 mM, black, n = 10) or NSF_p_ (1 mM, red, n = 9). Traces on the left and middle are single traces, whereas traces on the right are normalized, averaged traces. (D) Similar to panel C, but with SNAP_sp_ (1 mM, black, n = 7) or SNAP_p_ (1 mM, red, n = 8). (E) The Rate_decay_n_, ΔCm_35s_, ΔCm_peak_, and QICa induced by depol_20ms_ at 4 - 10 min after break in with a pipette containing the control solution (Ctrl, n = 11), ATPγS (4 mM, n = 6), NEM (1 mM, n = 12), NSF_mp_ (1 mM, n = 10), NSF_p_ (1 mM, n = 9), SNAP_sp_ (1 mM, n = 7) or SNAP_p_ (1 mM, n = 8). (F-J) Similar arrangement as panel A-E (including the calyx number), respectively, except that the stimulus was depol_20msX10_ and the ΔCm_35s_ was replaced with ΔCm_50s_ (E). The Cm decay in the Ctrl trace in B was fit bi-exponentially with τ of 1.4 s (weight: 29%) and 18.3 s, respectively (gray).

#### Quantifying slow and fast endocytosis in control

We first analyzed data during 4 - 10 min after whole-cell break-in. In control, we induced slow and fast endocytosis with 1 and 10 pulses of 20 ms depolarization (from -80 to +10 mV, if not mentioned otherwise) at 10 Hz, called depol_20ms_ (Fig. 1A) and depol_20msX10_ (Fig. 1F), respectively^8–10^ (Wu et al., 2005; Wu et al., 2009). A depol_20ms_ induced a capacitance jump (ΔCm_peak_) of 531 ± 55 fF (n = 11), followed by a slow decay with a τ of 10.2 ± 0.9 s (n = 11) and an initial decay rate (Rate_decay_, see Methods) of 56 ± 9 fF/s (n = 11, e.g., Fig. 1A). A depol_20msX10_ induced a ΔCm_peak_ of 1565 ± 115 fF (n = 11), followed by a bi-exponential decay with a rapid τ of 1.6 ± 0.3 s (amplitude: 31 ± 4%) and a slow τ of 15.5 ± 1.4 s (n = 11, e.g., Fig. 1F). The Rate_decay_ after depol_20msX10_ was 301 ± 55 fF/s (n = 11, e.g., Fig. 2B, left), which reflected mostly (>80%) the rapid component of endocytosis as demonstrated previously^8–10^ (Wu et al., 2005; Wu et al., 2009). When NSF_mp_ or SNAP_sp_ was included, which served as the control for NSF_p_ or SNAP_p_, it did not significantly affect all results described above (Figs. 1C-E, 1H-J). These results were similar to our previous reports^8–10^.

**Figure 2.**
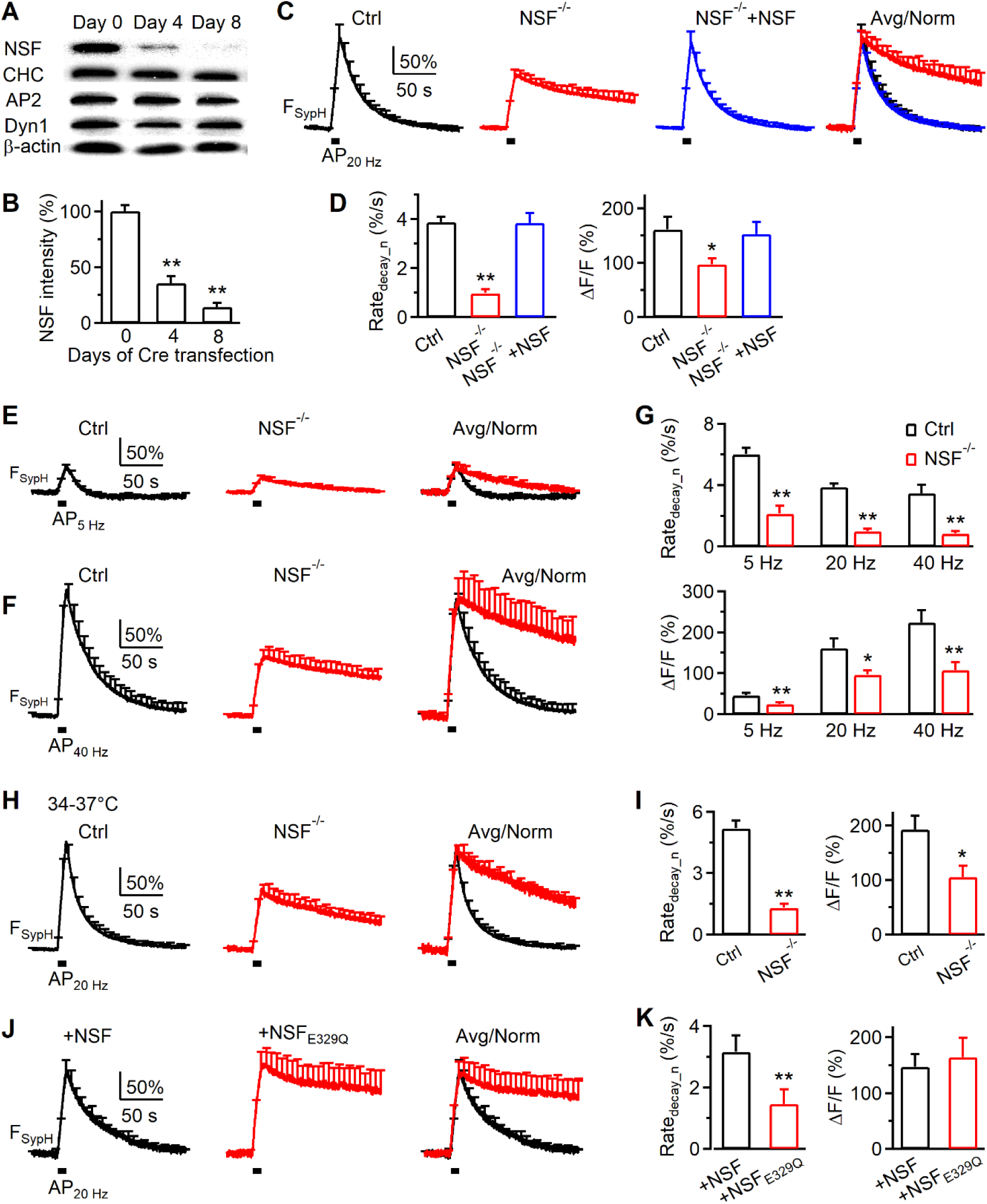
NSF knockout inhibits endocytosis at hippocampal synapses. (A-B) Sampled western blot of NSF, clathrin heavy chain (CHC), adaptor protein 2 α subunit (AP2), dynamin 1 (Dyn 1), and β-actin at day 0, day 4, and day 8 after Cre-4-OHT treatment to the *NSF^LoxP/LoxP^* hippocampal culture. (B) The NSF intensity (mean + s.e.m. from 8 cultures) recorded from the western blot at day 0, day 4, and day 8 after Cre-4-OHT treatment to the *NSF^LoxP/LoxP^* hippocampal culture. (C-D) The traces (C), Rate_decay_n_ (D) and ΔF/F (D) of SypH fluorescence (F_SypH_) changes (mean + s.e.m.) induced by Train_20Hz_ in control (n = 20 experiments, wildtype, black), NSF^-/-^ hippocampal boutons (n = 19 experiments, red), and NSF^-/-^ boutons overexpressed with wildtype NSF (NSF^-/-^+NSF; blue; containing EBFP2 for recognition, n = 23) at 22-24 °C. Traces are also normalized and overlapped to show the block of the F_SypH_ decay (C, right). *, p < 0.05; **, p < 0.01 (t test). (E-G) The traces (E, F), Rate_decay_n_ (G) and ΔF/F (G) of F_SypH_ changes (mean + s.e.m.) induced by a 10 s AP train at 5 Hz (E, G) or 40 Hz (F, G) in control (5 Hz: 11 experiments; 40 Hz: 11 experiments; black) and in NSF^-/-^ hippocampal boutons (5 Hz: 10 experiments; 40 Hz: 10 experiments; red) at 22-24 °C. Traces are also normalized and overlapped to show the block of the F_SypH_ decay (E-F, right). Rate_decay_n_ and ΔF/F induced by AP_20Hz_ are also included in panel G for comparison. (H-I) The traces (H), Rate_decay_n_ (I) and ΔF/F (K) of F_SypH_ changes (mean + s.e.m.) induced by Train_20Hz_ in control (n = 19 experiments, wildtype, black) and NSF^-/-^ hippocampal boutons (n = 9 experiments, red) at 34-37 °C. Traces are also normalized and overlapped to show the block of the F_SypH_ decay (H, right). (J-K) The traces (J), Rate_decay_n_ (K) and ΔF/F (K) of F_SypH_ changes (mean + s.e.m.) induced by Train_20Hz_ in wildtype cultures expressed with wildtype NSF (n = 14 experiments, +NSF, black) or NSF_E329Q_ (n = 8 experiments, +NSF_E329Q_) at 22-24 °C. Traces are also normalized and overlapped to show the block of the F_SypH_ decay (J, right).

When the ΔCm_peak_ was normalized to 1, the normalized Rate_decay_ (Rate_decay_n_) after depol_20ms_ was 0.09 ± 0.01/s (n = 11), meaning 9% of ΔCm_peak_ was retrieved at the first second after stimulation. At 35 s after depol_20ms_, the remaining ΔCm (ΔCm_35s_) was 4 ± 3% (n = 11) of the ΔCm_peak_, indicating complete endocytosis (Fig. 1A). The Rate_decay_n_ after depol_20msX10_ was 0.12 ± 0.01/s (n = 11), which largely reflected the normalized rapid endocytosis rate^8–10^. At 50 s after depol_20msX10_, the remaining ΔCm (ΔCm_50s_) was 3 ± 2% (n = 11) of the ΔCm_peak_, indicating complete endocytosis (Fig. 1F). Throughout the study, we compared these normalized values (Rate_decay_n_ and ΔCm_35s_ or ΔCm_50s_) in control and in the presence of drugs. We did not compare τ, because τ was often too long to estimate in the presence of blockers (Fig. 1). We did not compare Rate_decay_, because it is influenced by the ΔCm_peak_, which was often reduced by the tested blockers.

#### Inhibition of NSF inhibits both slow and fast endocytosis

Compared to the corresponding control, four NSF blockers at 4 - 10 min after break in substantially reduced the Rate_decay_n_ induced by either depol_20ms_ or depol_20msX10_ to < 40%, increased ΔCm_35s_ after depol_20ms_ or ΔCm_50s_ after depol_20msX10_ to > 25% (of ΔCm_peak_) (Fig. 1B-E, 1G-J), but did not affect the calcium current charge (QICa) or amplitude (Figs. 1E, 2E, Fig. S1). Consequently, these four types of blockers substantially prolonged the normalized, averaged capacitance decay (Figs. 1B-D, 1G-I). These results suggest that NSF is involved in mediating both slow and fast endocytosis.

#### Ratedecay_n reduction is independent of the ΔCmpeak reduction

Except for ATPγS and 0 ATP, other blockers reduced ΔCm_peak_ to a value >60% of control (Figs. 1B-E, 1G-J). Four sets of evidence suggest that block of ΔCm_peak_ did not contribute to the decrease of Rate_decay_n_ or the increase of ΔCm_35s_ or ΔCm_50s_. First, Rate_decay_n_ was normalized to the ΔCm_peak_, which normalized the contribution of the ΔCm_peak_ decrease on Rate_decay_. Second, in control, increase of exocytosis is accompanied by an increase of the endocytosis τ when endocytic capacity is saturated^8,11–14^, or no change of τ when endocytic capacity is not saturated^15^. Thus, as the exocytosis amount was decreased by NSF blockers, the endocytosis τ should decrease or remain unchanged, which means that Rate_decay_n_ should increase or remain unchanged if endocytosis is not inhibited. The observed decrease of the Rate_decay_n_ (Figs. 1-2) is therefore not caused by reduction of exocytosis itself, but by inhibition of endocytosis. Third, ATPγS blocked endocytosis, but not ΔCm_peak_ or QICa within 4 min after break in (Fig. 1B, 1G), indicating that endocytosis block can be independent of the ΔCm_peak_. At later dialysis time points, ATPγS reduced ΔCm_peak_ and QICa. We did not analyze these data because the QICa decrease may complicate analysis of the Rate ^9^. Fourth, at 2 - 4 min after whole-cell break in, during which the exocytosis block was minimal, NSF_p_ and SNAP_p_ did not decrease the ΔCm_peak_, but still significantly reduced the Rate_decay_n_ induced by depol_20ms_ (Fig. S2A-C) or depol_20msX10_ (Fig. S2D-F).

### NSF and its SNARE disassembly activity are required for endocytosis at hippocampal synapses

The specificity of pharmacological blockers is a common concern in pharmacology experiments. We addressed this concern by 1) using four different NSF blockers in the calyx of Held, all of which generate the consensus inhibition of endocytosis (Fig. 1), and 2) knocking out or knocking down NSF genes as described below.

#### NSF conditional knockout mouse generation and gene deletion in culture synapses

To block NSF specifically, we generated NSF conditional knockout (*NSF^LoxP/LoxP^*) mice by floxing *NSF* Exon 6 and Exon 7 (Fig. S3). In hippocampal neurons cultured from *NSF^LoxP/LoxP^* mice, we deleted *NSF* by treating the culture with Cre-4-OHT or by Cre transfection (with mCherry for recognition)^16^. Western blot showed that Cre-4-OHT treatment progressively reduced NSF to below 20% in 8 days, but does not affect other endocytic proteins (dynamin, clathrin, adaptor protein 2) (Fig. 2A-B). Similar reduction was obtained at 8 days after Cre transfection, as detected with immunostaining (Fig. S4). Since these two methods resulted in similar NSF reduction, we grouped them together as the NSF^-/-^ culture, and their corresponding data were thus accordingly.

#### SynaptopHluorin imaging of endocytosis

To record endocytosis, we transfected pH-sensitive synaptopHluorin (SypH) to the NSF^-/-^ hippocampal culture and imaged SypH fluorescence (F_SypH_) from boutons at room temperature, if not mentioned otherwise^11,17^. A train of action potential stimulation at 20 Hz for 10 s (AP_20Hz_) induced a F_SypH_ increase and decrease, reflecting exocytosis and endocytosis, respectively (Fig. 2C). In control (*NSF^LoxP/LoxP^* culture), the peak F_SypH_ increase over the baseline (ΔF/F) is 161.0 ± 23.4% (Fig. 2D); F_SypH_ decay is mono-exponential with an initial decay rate (Rate_decay_n_) of 3.9 ± 0.2%/s (n = 20 experiments, each experiment contained ∼10 - 30 boutons; Fig. 2D), where the rate was normalized to ΔF/F.

In *NSF*^-/-^ cultures, ΔF/F induced by AP_20Hz_ decreased ΔF/F to ∼60% of control, and reduced Rate_decay_n_ to about 25% of control (Fig. 2C-D, n = 19 experiments). Both ΔF/F and Rate_decay_n_ were rescued to the control level by transfection of WT NSF to the *NSF*^-/-^ culture (Figure 2C-D, 20 experiments). These results suggest that NSF is required for mediating endocytosis after 10 Hz nerve firing.

Similar inhibition of both ΔF/F and Rate_decay_n_ was observed after 5 or 40 Hz action potential stimulation for 10 s (Fig. 2E-G), or AP_20Hz_ at physiological temperature (^34∼37°C, Fig. 2H-I^). These results suggest that NSF is required for endocytosis regardless of the stimulation frequency and temperature.

#### ATP hydrolysis is required for endocytosis

The ATPase NSF hydrolyses ATP to disassemble the SNARE complex, which was blocked by a NSF mutant E329Q (NSF_E329Q_)^18,19^. To determine whether NSF’s ATPase function in disassembling the SNARE complex is needed for endocytosis, we overexpressed NSF or NSF_E329Q_ to WT hippocampal neurons. We found that AP_20Hz_-induced Rate_decay_n_ was ∼3.3%/s with NSF transfection (control), but was reduced to ∼44% of control with NSF_E329Q_ transfection (Fig. 2J-K), suggesting that ATPase-mediated SNARE complex disassembly is required for endocytosis at hippocampal synapses.

#### NSF involvement in bulk endocytosis at hippocampal synapses observed with EM

We performed electron microscopy (EM) to examine the ultrastructural changes in NSF^-/-^ hippocampal cultures at physiological temperature. Horseradish peroxidase (HRP, 5 mg/ml) was added in bath for assay of vesicular uptake. At rest, HRP-positive [HRP(+)] vesicles were minimal; most vesicles were HRP-negative [HRP(-)] (Fig. 3A); the number of HRP(+) vesicles in boutons was similar in Ctrl and NSF^-/-^ cultures. To examine endocytosis, we applied 90 mM KCl with HRP for 1.5 min, and fixed samples at 0, 3 and 10 min after KCl/HRP application. In Ctrl boutons, compared with the resting condition, HRP(+) vesicles increased from time 0 to 10 min after KCl, reflecting vesicle endocytosis (Fig. 3A-B) as previously shown^20,21^. Compared with Ctrl boutons, HRP(+) vesicles were significantly reduced at each time point after KCl application in NSF^-/-^ boutons (Fig. 3A-B), suggesting inhibition of endocytosis of regular vesicles.

**Figure 3.**
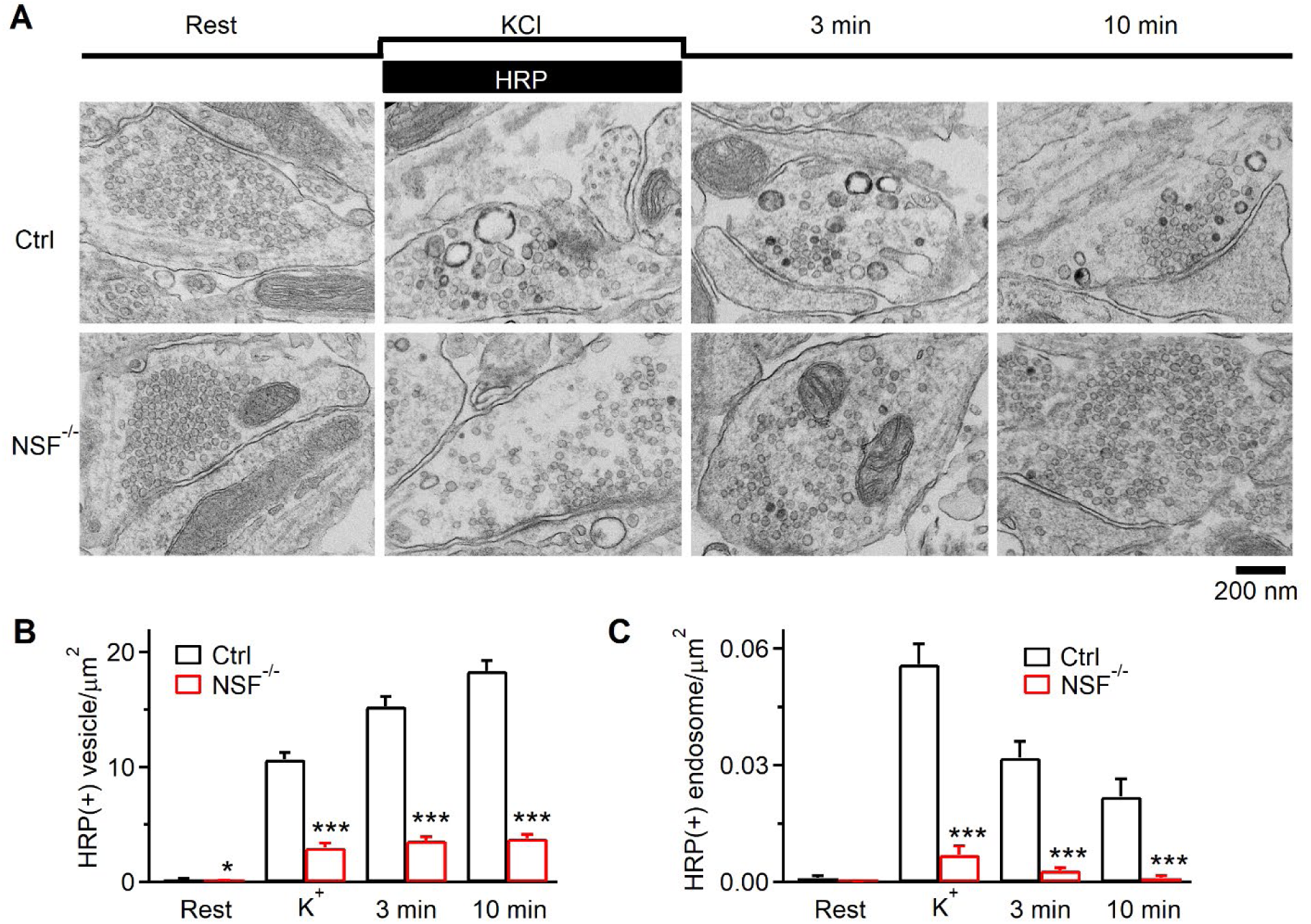
NSF knock-out affects endocytosis examined with EM at hippocampal synapses. (A) EM images of WT and NSF^-/-^ hippocampal boutons at rest (Rest) and at 0 min (KCl), 3 min, and 10 min after 1.5 min 90 mm KCl application. For Rest, HRP was included for 1.5 min; for KCl application, HRP was included only during KCl application (see labels). (B-C) Number of HRP(+) vesicles (B) and the bulk endosome area (C) per square micrometer of synaptic cross-section are plotted versus the time before (Rest) and at 0 min (K+), 3 min, and 10 min after the end of KCl application in WT and NSF^-/-^ hippocampal cultures (mean + SEM, each group was from 120–122 synaptic profiles from 18 mice (6 mice per experiment, 3 experiments). The temperature before fixation was 37°C. *: p<0.05; ***: p<0.001, t-test.

In Ctrl boutons, we observed HRP(+) bulk endosomes (Fig. 3A), defined as vesicles with a diameter 80 nm or with a cross section area more than that of a 80 nm vesicle (∼0.005 µm^2^). Bulk endosome area increased at time 0, then decreased at 3 and 10 min (Fig. 3A, C), suggesting generation of bulk endosomes and subsequent conversion to vesicles as previously shown^10,20^. Similar trends were observed in NSF^-/-^ cultures, but at a significantly lower level (Fig. 3A, C), suggesting inhibition of bulk endocytosis. Thus, EM results reveal the involvement of NSF in regular vesicle endocytosis and bulk endocytosis at hippocampal synapses.

### NSF is essential for pore closure of preformed and fusion-generated Ω-profiles in chromaffin cells

We showed NSF involvement in slow, fast, and bulk endocytosis at synapses (Figs. 1-3). Since these different endocytic modes are mediated primarily by the pore closure of preformed Ω-profiles (pre-Ω, formed before depolarization) and fusion-generated Ω-profiles (fs-Ω) in chromaffin cells^22–24^, we determined whether NSF involvement in endocytosis is due to its role in closing pre-Ω and fs-Ω’s pore in chromaffin cells in the following.

#### Methods for imaging pore closure

We have developed imaging methods to detect pre-Ω and fs-Ω pore closure in live adrenal chromaffin cells^22–24^. Pre-Ω (∼200-1500 nm in diameter) could be generated from 1) the endocytic flat-to-Ω-shape transition, including bulk endocytosis that produces vesicles larger than fusing vesicles^24,25^, and 2) dense-core vesicle fusion, some of which could maintain the Ω-shape for a long time^24^. Fs-Ω is from fusion of dense-core vesicles^23^ with a diameter of ∼360 nm (range: 200-700 nm)^26,27^. We used mNeonGreen attached to phospholipase C δPH domain (PH_G_, overexpressed, binds to PI(4,5)P_2_) to label the plasma membrane (PM), Atto 655 (A655, 30 μM in bath; or Atto 532) to fill Ω-profiles, and fluorescent false neurotransmitter FFN511 (or FFN206) pre-loaded into vesicles to measure release (Fig. S5; Fig. 4A-C)^16,24^. At the bottom plasma membrane of resting cells, XY-plane confocal microscopy observed FFN511-containing vesicle spots and preformed PH_G_ spots and rings overlapped with A655, but not FFN511 spots (termed pre-spot, Fig. S5). Pre-spots were mostly pre-Ω as observed at the XZ-plane with confocal or stimulated emission depletion (STED) microscopy (e.g., Fig. S5, for detail, see Ref. ^24^).

**Figure 4.**
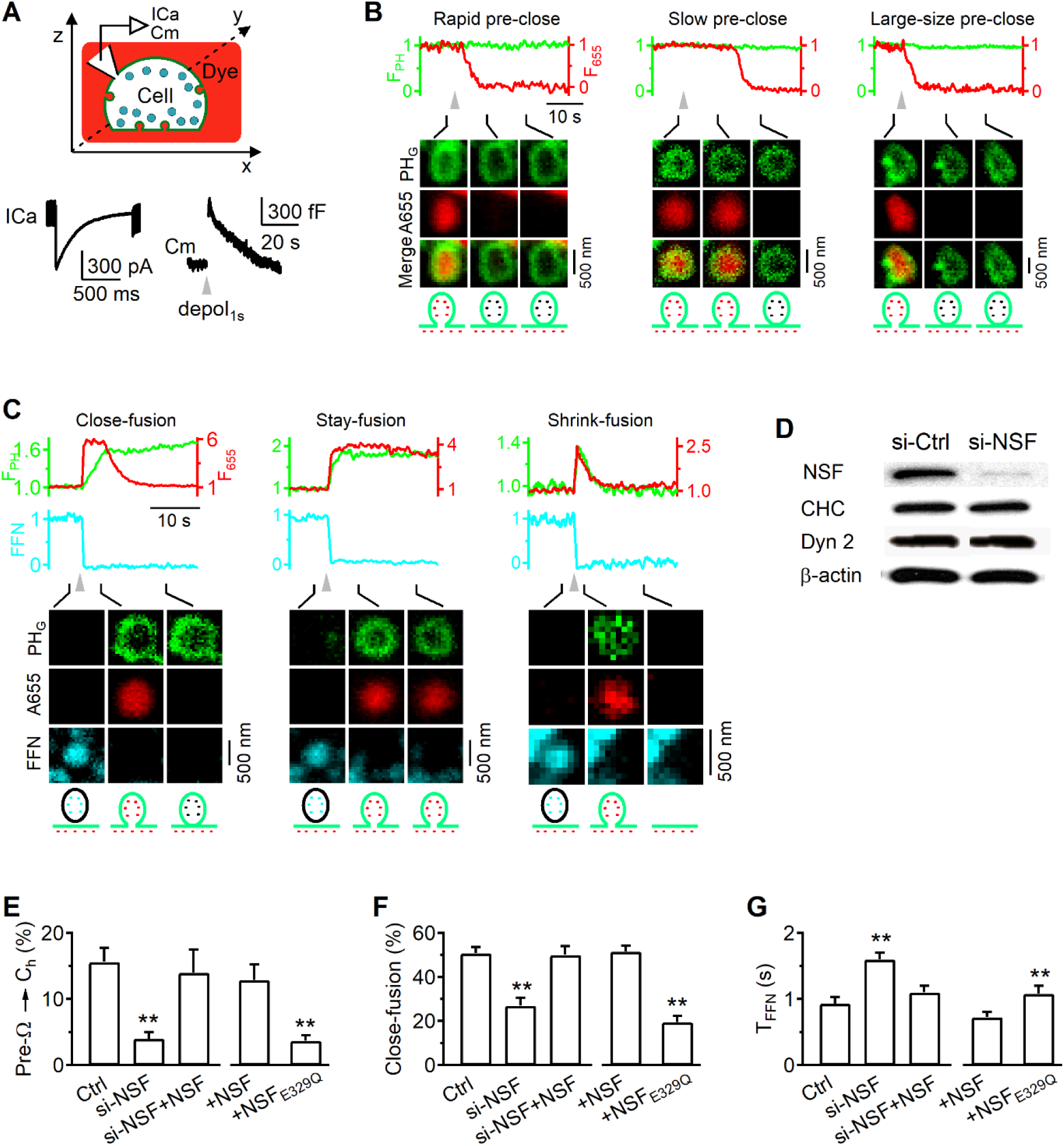
NSF is essential for mediating pre-Ω and fs-Ω pore closure in chromaffin cells. (A) Upper: setup drawing. The cell membrane, bath and vesicles are labelled with PH_G_ (green), A655 (red) and FFN511 (yellow), respectively. ICa and Cm (capacitance) are recorded via a whole-cell pipette. Lower: Sampled ICa and Cm changes induced by depol_1s_. (B) PH_G_ fluorescence (F_PH_), A655 fluorescence (F_655_) and sampled confocal images showing depol_1s_-induced (triangle) rapid (left), slow (middle) or large-size (right) pre-spot pore closure (pre-close) in chromaffin cells. F_PH_ and F_655_ were normalized to the baseline. (C) F_PH_, F_655_, FFN511 fluorescence (F_FFN_), and confocal images showing close-fusion, stay- or shrink-fusion. (D) Western blot of NSF, clathrin heavy chain (CHC), dynamin 2 (Dyn 2), and β-actin in chromaffin cell cultures transfected with si-Ctrl or si-NSF. (E-G) The percentage of pre-spots undergoing depol_1s_-induced pre-close (E), the percentage of fusing vesicles undergoing close-fusion (F), and the 20-80% decay time of F_FFN_ (G, indicating release time) in control (Ctrl, n = 25 cells), si-NSF transfection (n = 22 cells), si-NSF transfection plus wildtype NSF overexpression (si-NSF+NSF, 15 cells), control cells overexpressed with wildtype NSF (+NSF, 22 cells), or control cells overexpressed with NSF_E392Q_ (+NSF_E932Q_, 24 cells). *: p < 0.05; **: p < 0.01(t test, compared to control).

A whole-cell 1-s depolarization (−80 to +10 mV, depol_1s_) induced ICa, capacitance changes reflecting exo-endocytosis (Fig. 4A), pre-spot closure (pre-close, Fig. 4B), and fusion spots observed with confocal microscopy (Fig. 4C; cell-bottom, XY-plane imaging every 40-80 ms)^22–24^. Pre-close was detected as A655 fluorescence (F_655_, strongly excited) dimming while PH_G_ fluorescence (F_PH_, weakly excited) sustained or dimmed with a delay (Fig. 4B)^24^. This method detected pore closure of pre-Ω that was impermeable to H^+^ and OH^-^, mediated by dynamin, and observed directly with STED imaging^22–24,28^. Pre-Ω closure forms ∼200-1500 nm vesicles^24^, with ∼17% in the 600 - 1500 nm range (e.g., Fig. 4B, right) that can be attributed to bulk endocytosis^24^.

Fusion spots were detected as a sudden appearance of PH_G_ and A655 spots while FFN511 spot fluorescence (F_FFN_) decayed, due to the diffusion of PH_G_/A655 from the PM/bath to the fs-Ω and release of FFN511 from the fs-Ω (Fig. 4C). Three fusion modes were observed (see Methods for more detail): 1) close-fusion (kiss-and-run) – fs-Ω pore closure was detected similarly to pre-close: as F_655_ dimming while F_PH_ was sustained or decayed later (Fig. 4C, left)^22–24,28^; 2) stay-fusion – a sustained fs-Ω was detected as persistent PH_G_/A655 spots with sustained F_655_ and F_PH_ (Fig. 4C, middle); 3) shrink-fusion – fs-Ω shrinking was detected as parallel decreases of spot-size with F_655_ and F_PH_ (Fig. 4C, right)^23,28,29^. STED imaging directly observed these modes (for detail, see Refs. ^23,29^).

#### NSF and its ATPase activity are required for pre-Ω and fs-Ω pore closure

NSF siRNA (si-NSF) transfection substantially reduced NSF without affecting key endocytic protein dynamin and clathrin (Fig. 4D). si-NSF blocked depol_1s_-induced pre-close (Fig. 4E) and close-fusion (Fig. 4F), but increased the FFN511 20-80% decay time that reflects the release time course (T_FFN_ – release time, Fig. 4G). NSF overexpression in si-NSF-transfected cells rescued pre-close (Fig. 4E), close-fusion (Fig. 4F), and T_FFN_ (Fig. 4G) to the control level. These results suggest that NSF is required for pre-close and close-fusion, and controls the time course of vesicular content release.

Overexpression of NSF_E329Q_, a mutant unable to disassemble SNARE complexes^18,19^, inhibited depol_1s_-induced pre-close (Fig. 4E) and close-fusion (Fig. 4F), but increased T_FFN_ (Fig. 4G), suggesting that SNARE disassembly by NSF is required for closing both pre-Ω and fs-Ω’s pore, and controlling content release time course.

### NSF mediates slow, fast, ultrafast, and overshoot endocytosis by closing pre-Ω/fs-Ω’s pore in chromaffin cells

Since endocytosis in chromaffin cells is primarily due to pre-Ω/fs-Ω’s pore closure rather than the generally believed flat-to-round endocytic transformation^16,24^, our finding of NSF involvement in pre-Ω/fs-Ω pore closure predicts NSF involvement in endocytosis. We verified this prediction by examining how inhibition of NSF affects depol_1s_-induced capacitance decay (after the jump) that reflects endocytosis. The following two sets of results suggest that NSF and its SNARE-disassembly function are required for endocytosis in chromaffin cells. First, si-NSF transfection inhibited the Cm-decay-indicated endocytosis, but not ICa; and the inhibition was rescued by overexpression of wild-type NSF (Fig. 5A). Second, overexpression of NSF_E329Q_ inhibited the Cm-decay-indicated endocytosis as compared to overexpression of wildtype NSF (Fig. 5B).

**Figure 5.**
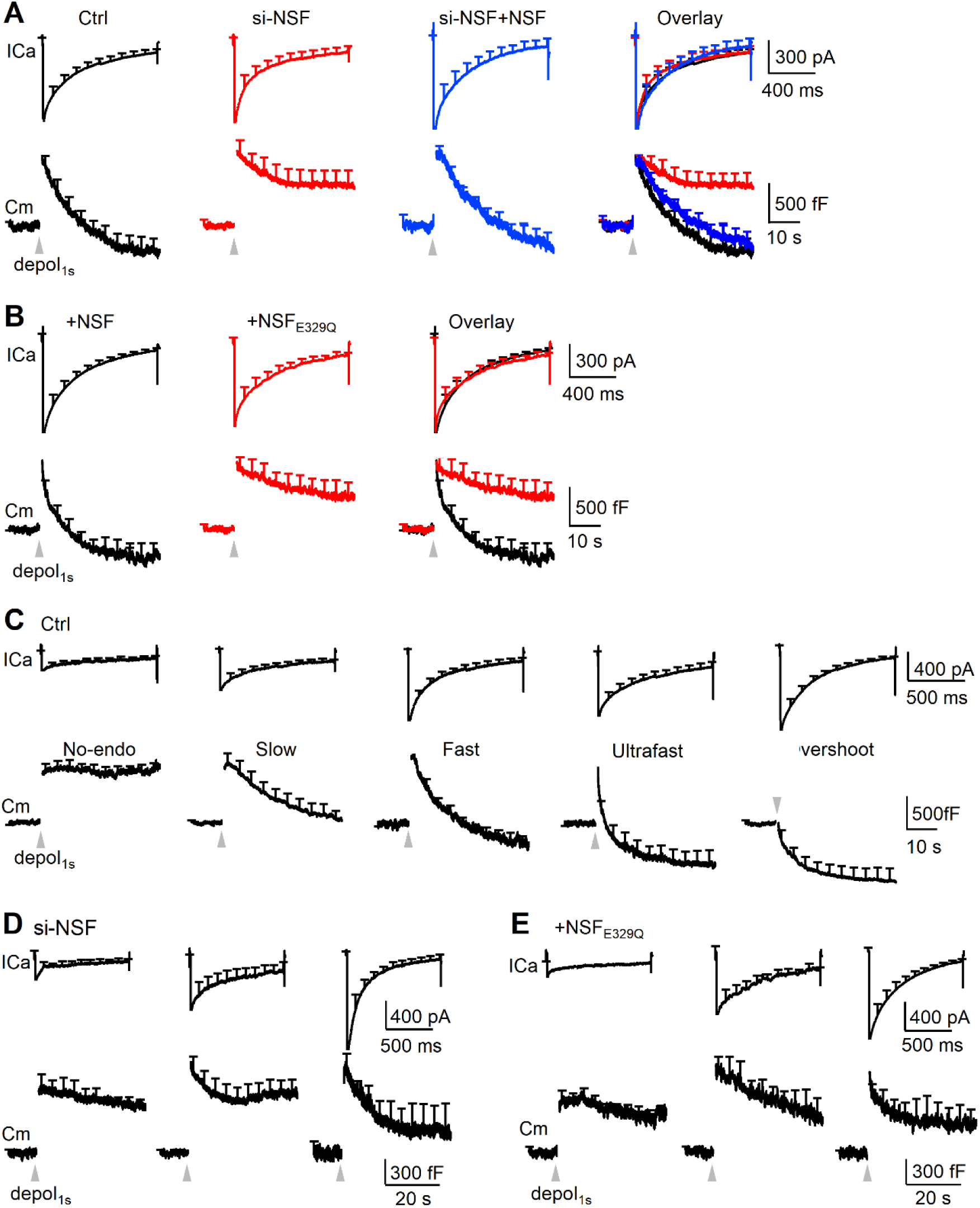
NSF is essential for mediating slow, fast, ultrafast, and overshoot endocytosis in chromaffin cells. (A) Depol_1s_-induced ICa (mean + s.e.m., upper) and Cm (mean + s.e.m., lower) in chromaffin cells in control (Ctrl, 25 cells), si-NSF transfection (si-NSF, 22 cells), and si-NSF transfection plus wildtype NSF overexpression (si-NSF+NSF, 15 cells). Traces are also merged in the right. (B) Depol_1s_-induced ICa (mean + s.e.m., upper) and Cm (mean + s.e.m., lower) in chromaffin cells overexpressed with wildtype NSF (+NSF, 22 cells) or NSF_E392Q_ (+NSF_E392Q_, 24 cells). Traces are also merged in the right. (C) Mean ICa (mean + s.e.m., upper) and Cm (mean + s.e.m., lower) induced by depol_1s_ (gray triangle) in five groups of chromaffin cells (from left to right): Group_no-endo_ (decay < 30% ΔCm, 6 cells), Group_slow_ (endocytic τ > 6 s, 6 cells), Group_fast_ (τ: 0.6 – 6 s, 5 cells), Group_ultrafast_ (τ < 0.6 s, 4 cells) and Group_overshoot_ (decay > 130% ΔCm, 4 cells) in control chromaffin cells.(D-E) Depol_1s_-induced ICa (mean + s.e.m., upper) and Cm (mean + s.e.m., lower) in chromaffin cells with ICa of 160 – 360 pA (left), 400 – 900 pA (middle), and 1000 – 1800 pA (right) in two conditions: (D) si-NSF transfection (left: 8 cells; middle: 7 cells; right: 7 cells); (E) NSF_E392Q_ overexpression (left: 9 cells; middle: 8 cells; right: 7 cells).

To determine which mode(s) of endocytosis NSF is involved in, we first described five distinct modes of endocytosis previously characterized in chromaffin cells{Shin, 2021 #2275}{Wei, 2024 #2525}. Five modes of endocytosis were revealed when chromaffin cells were divided into five groups based on the decay of the whole-cell capacitance (Cm) after the jump induced by depol_1s_: 1) no-endocytosis (Group_no-endo_, decay <30% ΔCm), 2) slow endocytosis (Group_slow_, endocytic τ > 6 s), 3) fast endocytosis (Group_fast_, τ: 0.6 - 6 s), 4) ultrafast endocytosis (Group_ultrafast_, τ < 0.6 s), and 5) overshoot endocytosis (Group_overshoot_, decay > 130% ΔCm, Fig. 5C; see Ref. ^24^ for detail). Calcium influx triggers endocytosis in each of these five groups: larger ICa induces faster and larger amplitude of endocytosis (Fig. 5C), whereas strontium abolishes endocytosis (see Ref. ^24^ for detail).

si-NSF transfection or NSF_E329Q_ overexpression inhibited the Cm-decay-indicated endocytosis in all three cell groups divided based on the ICa amplitude (Fig. 5D-E), indicating inhibition of endocytosis throughout the entire ICa range that generates five distinct endocytic modes in control. These results suggest that NSF and its SNARE complex disassembly function are required to drive slow, fast, ultrafast, and overshoot endocytosis observed in control. We concluded that NSF and its SNARE complex disassembly function contribute to mediating diverse modes of endocytosis by participating in closing the pre-Ω and fs-Ω’s pore.

## Discussion

We demonstrated that NSF inhibition —via pharmacological blockers, gene knockout, and knockdown— in the calyx of Held nerve terminal, hippocampal boutons, and neuroendocrine chromaffin cells impaired multiple modes of endocytosis, including slow, fast, ultrafast, overshoot, and bulk endocytosis (Figs. 1-3, 5). These effects were consistently observed across different experimental techniques, including capacitance measurements, synaptopHluorin imaging, and electron microscopy, and were reproducible with different NSF inhibition strategies across multiple cell types. Notably, inhibition of NSF’s ATPase activity, which is essential for SNARE complex disassembly, replicated these findings (Figs. 1, 2, 5). In chromaffin cells, we further demonstrated that these effects result from inhibition of pre-Ω/fs-Ω pore closure (Fig. 4) that is responsible for generating the full spectrum of endocytic modes described above^16,24^. Together, these results establish NSF as a key molecular component of the endocytic machinery governing multiple endocytic modes and kiss-and-run fusion. Specifically, NSF emerges as a central regulator of fission and fusion pore closure in diverse endocytic pathways that recycle exocytosed vesicles. We propose revising existing models of membrane fission, kiss-and-run fusion, endocytosis, and vesicle recycling to incorporate NSF as a critical molecular player.

While finding the involvement of the core ‘exocytosis’ protein NSF in endocytosis is apparently surprising, it is consistent with a series of studies showing that other core exocytosis proteins, including SNAP-25, syntaxin, VAMP2, and synaptotagmin 1, are involved in synaptic vesicle endocytosis at calyx-type and hippocampal synapses^30–36^. The dual roles of these core ‘exocytosis’ proteins in both exo- and endocytosis suggest that they may play an important role in coupling exocytosis to endocytosis, which recycles vesicles and maintains the exocytosis capacity and membrane homeostasis of release sites^37,38^.

The dual exo-endocytosis role also suggests that exo- and endocytosis may take place at the same location or nearby. Indeed, we demonstrated that NSF and its SNARE-disassembly function are required to close the pore of fs-Ω (fusion pore) and pre-Ω, thereby mediating endocytosis in chromaffin cells (Figs. 4, 5). Given that pre-Ω could be generated by fusing vesicles that maintain a Ω-shape, NSF and its SNARE-disassembly function may mediate fusion pore closure and thus endocytosis via the kiss-and-run (close-fusion) and kiss-and-stay fusion (stay-fusion). Since pre-Ω could also be generated from endocytic flat-to-Ω shape transition^24,39^, NSF and its SNARE-disassembly function may be required for pore closure during classical endocytic flat-to-round vesicle formation.

How is NSF involved in pore closure? A recent study shows that SNARE disassembly by NSF is essential for closing the fusion pore formed in the *in vitro* SNARE-reconstituted nanodisk^40^. The SNARE complex formation is required to assemble a fusion pore, whereas the SNARE complex disassembly by NSF has been suggested to disassemble the fusion pore, resulting in the fusion pore closure^40^. This *in vitro* finding offers a mechanistic explanation for why NSF and its SNARE-disassembly function are required to mediate fusion pore closure in live chromaffin cells.

This explanation seems difficult to account for the requirement of NSF and its SNARE-complex disassembly function in closing the pre-Ω’s pore. It might be possible that after fusion, the SNARE complex is sorted to the endocytic site around the pre-Ω’s pore region, which may present a physical barrier that may prevent dynamin from accessing the pore. Disassembly of the SNARE complex at the pre-Ω by NSF might thus facilitate pre-Ω pore closure, explaining why NSF is needed for the pre-Ω pore closure. While beyond the scope of the present work, it is interesting to determine how NSF is involved in pore closure in the future.

Our findings provide an explanation for a long-standing observation that endocytosis requires energies from not only GTP hydrolysis^14,41^, but also ATP hydrolysis^42^ – NSF hydrolyzes ATP to disassemble the SNARE complex. Through its ATPase activity that disassembles the SNARE complex, NSF 1) mediates diverse modes of endocytosis, as shown here, 2) regulates the trafficking of neurotransmitter receptors, including AMPA receptors, GABA receptors, and dopamine receptors, and 3) contributes to generating synaptic plasticity in the nervous system. across different types of synapses^6^.

The dysfunction of NSF-mediated trafficking of these receptors or NSF mutations is associated with several neurological disorders, such as Alzheimer’s disease and epilepsy^6^. Our finding that NSF mediates diverse endocytic modes by closing pre-Ω and fs-Ω’s pore offers a novel mechanism accounting for these physiological and pathological roles of NSF.

## Acknowledgements

We thank Jianhua Xu for the strong support of calyx experiments, Susan Cheng and Virginia Crocker for EM technical support, Dr. Gero Miesenböck (University of Oxford, Oxford, UK) for providing us with the synaptopHluorin plasmid, and Dr. Yongling Zhu for synaptophysin-pHluorin2X plasmid. This work was supported by NINDS Research Program (ZIA NS003009-15 and ZIA NS003105-10 to L.G.W.). The contributions of the NIH author(s) were made as part of their official duties as NIH federal employees, are in compliance with agency policy requirements, and are considered Works of the United States Government. However, the findings and conclusions presented in this paper are those of the author(s) and do not necessarily reflect the views of the NIH or the U.S. Department of Health and Human Services.

## Author contributions

X.S.W. performed imaging experiments in chromaffin cells and electrophysiology experiments; T.S., S.B., and S.L. performed imaging experiments in hippocampal synapses; S.B. performed EM experiments; Z.Z. L.W., X.W., M.M., S.H. participated in experiments, L.G. generated NSF conditional knockout mice; X.S.W. S.B. and L.G.W. wrote the manuscript with help from all authors. L.G.W. wrote the manuscript and supervised the project.

## Methods and Materials

### Slice preparation, capacitance recordings and solutions

Slice preparation and capacitance recordings were similar as previously described^9,12,43,44^. Briefly, parasagittal brainstem slices (200 μm thick) containing the medial nucleus of the trapezoid body were prepared from 7 - 10 days old male or female Wistar rats using a vibratome. Whole-cell capacitance measurements were made with the EPC-9 amplifier together with the software lock-in amplifier (PULSE, HEKA, Lambrecht, Germany) that implements Lindau-Neher’s technique. The frequency of the sinusoidal stimulus was 1000 Hz and the peak-to-peak voltage of the sine wave was ≤60 mV. We pharmacologically isolated presynaptic Ca^2+^ currents with a bath solution (∼22 - 24 °C) containing (in mM): 105 NaCl, 20 TEA-Cl, 2.5 KCl, 1 MgCl_2_, 2 CaCl_2_, 25 NaHCO_3_, 1.25 NaH_2_PO_4_, 25 glucose, 0.4 ascorbic acid, 3 *myo*-inositol, 2 sodium pyruvate, 0.001 tetrodotoxin (TTX), 0.1 3,4-diaminopyridine, 300 - 310 mOsm, pH 7.4 when bubbled with 95% O_2_ and 5% CO_2_. The presynaptic pipette contained (in mM): 125 Cs-gluconate, 20 CsCl, 4 MgATP, 10 Na_2_-phosphocreatine, 0.3 GTP, 10 HEPES, 0.05 BAPTA, 310 - 320 mOsm, pH 7.2, adjusted with CsOH.

NSF peptide (TGKTLIARKIGTMLNAREPK), mutated NSF peptide (TGKTLIARKIETMLNAREPK), SNAP peptide (QSFFSGLFGGSSKIEEACE), scrambled SNAP peptide (GFAESLFQSIEKESGFSCG) were purchased from the 21st Century Biochemicals, Inc. (Marlboro, MA, USA).

### Animals for hippocampal cells experiments

Animal care and use were carried out according to NIH guidelines and were approved by the NIH Animal Care and Use Committee (NINDS ASP 1170). Mouse housing conditions: temperature: 70-74 °F; humidity: 35-60%; light cycle: 6 AM-6 PM and dark cycle: 6 PM-6 AM. Wild-type (C57BL/6J) mice at P0 of either sex were used. *NSF*^Loxp^ mouse generation is produced by Dr. Lin Gan and described in Figure S3. *NSF*^Loxp/Loxp^ mice of either sex were obtained by heterozygous and homozygous breeding using standard mouse husbandry procedures. Mouse genotypes were determined by PCR.

### Mouse hippocampal culture and transfection

Mouse hippocampal culture was prepared as described previously^10,45^. Hippocampal CA1-CA3 regions from P0 wild-type mice were dissected, dissociated, and plated on Poly-D-lysine. Cells were maintained at 37°C in a 5% CO_2_ humidified incubator in a medium containing MEM, 0.5% glucose, 0.1 g/l bovine transferrin, 0.3 g/l glutamine, 10% fetal bovine serum, 2% B-27, and 3 µM cytosine β-D-arabinofuranoside. On 6-8 days after plating, neurons were transfected with plasmids using Lipofectamine LTX. Neurons were then maintained at 37 °C for 2 days before imaging.

Transfected plasmids included a plasmid containing synaptophysin-pHluroin (SypH) gifted by Dr. Yongling Zhu^46^ alone (control) or with a L309 plasmid containing Cre/mCherry. A nuclear localization sequence was tagged at the N-terminal of Cre, and cloned into L309 vector via BamHI and EcoRI sites. Accordingly, mCherry was expressed in the nucleus. For the rescue experiments (see Fig. 2), we transfected cDNA NSF plasmid (Novopro) along with SypH and Cre/mCherry. The cDNA encoding NSF was subcloned into EBFP2-C1 (Addgene #54665), and EBFP2 was used for us to recognize transfected cells.

The cDNA encoding NSF_E329Q_ (Genecopoeia) was subcloned into EBFP2-C1 (Addgene #54665) and EBFP2 was used for us to recognize transfected cells. For the rescue experiments, we transfected NSF plasmid along with SypH and NSF_E329Q_.

### Immunohistochemistry in hippocampal cultures

Cells were fixed with 4% paraformaldehyde, permeabilized with 0.3% Triton X-100, and subsequently incubated with primary and secondary antibodies. Primary antibodies were diluted in PBS containing 10% donkey serum and incubated with cells at 4 °C overnight. After several rinses in PBS, cells were incubated with fluorescence-conjugated donkey anti-mouse, anti-sheep, or anti-rabbit IgG (1:1000, Invitrogen) for 1 h at 22–24 °C. Primary antibodies included mouse anti-NSF (1:200, Abcam) and anti-TAU (1:200, Abcam). Imaging was similar to SypH imaging. mCherry fluorescence imaging was performed simultaneously to identify cells transfected with Cre/mCherry.

### SynaptopHluorin imaging in hippocampal neurons

Action potential was evoked by a 1 ms pulse (20 mA) through a platinum electrode. The bath solution contained (in mM): 119 NaCl, 2.5 KCl, 2 CaCl_2_, 2 MgCl_2_, 25 HEPES (buffered to pH 7.4), 30 glucose, 0.01 6-cyano-7-nitroquinoxaline-2, 3-dione (CNQX), and 0.05 D, L-2-amino-5-phosphonovaleric acid.

We heated the culture chamber using a temperature controller (TC344B, Warner Instruments, Hamden, CT). Imaging was performed after the culture was at 34-37°C for 15-30 min. The temperature was verified with another small thermometer (BAT-7001H, Physitemp Instruments, Clifton, NJ) in the chamber. SypH images were acquired at 10 Hz using Nikon A1 confocal microscope (Objective: 60x, 1.4 NA), and analyzed with Nikon software. All boutons showing fluorescence increases were analyzed (region of interest: 2 X 2 μm). Each data group was obtained from at least three batches of cultures.

F_SypH_ was normalized to the baseline F_SypH_ before stimulation (baseline F_SypH_ was normalized as 100%). Rate_decay_ (the initial rate of F_SypH_ decay) was measured from F_SypH_ in the first 4–10 s after stimulation. Each data group was obtained from ≥3 cultures; 1-3 experiments were taken from 1 culture; 10-30 boutons were used for each experiment.

### Electron microscope images, data collection and analysis of hippocampal neurons

Hippocampal cultures were fixed with 4% glutaraldehyde (freshly prepared, Electron microscopy sciences, Hatfield, PA) in 0.1 M Na-cacodylate buffer solution containing for at least 1 h at 22–24°C, and stored in 4°C refrigerator overnight. The next day, cultures were washed with 0.1 M cacodylate buffer, and treated with 1% OsO_4_ in cacodylate buffer for 1 h on ice, and 0.25% uranyl acetate in acetate buffer at pH 5.0 overnight at 4°C, dehydrated with ethanol, and embedded in epoxy resin. Thin sections were counterstained with uranyl acetate and lead citrate then examined in a JEOL200CX TEM. Images were collected with a CCD digital camera system (XR-100; AMT) at a primary magnification of 10,000 –20,000X. Synapses were selected based on the structural specialization including synaptic vesicle clustering, synaptic cleft and the postsynaptic density.

Data are presented as means ± SEM. The statistical test used was t test with equal variance, although t test with unequal variance gave the same result. For capacitance measurements at calyces, each group of data were from 4 –14 calyces, which were from 4 –14 mice of either sex (see figure legends for the number of calyces and mice for each group of data). For pHluorin imaging, each experiment included 20 –30 boutons showing fluorescence increase (region of interest: 2 µm x 2 µm). Approximately one to three experiments were taken from each culture. Each culture was from 3–5 mice. Each group of data was obtained from at least four batches of cultures (4 –12 cultures). For electron microscopy, synapses were selected based on the structural specialization, including synaptic vesicle clustering, synaptic cleft, and the postsynaptic density. Each group of data was taken from 100 –132 synaptic profiles from 4 –12 mice.

### Chromaffin cell culture and transfection

The primary bovine adrenal chromaffin cell culture has been descripted previously^22,28,47^. We purchased fresh adrenal glands (from 21 ̶ 27 months old bovines) from a local slaughterhouse (J. W. Treuth & Sons Inc., 328 Oella Ave, Catonsville, MD 21228; web site: https://www.jwtreuth.com). The glands were immersed in pre-chilled Locke’s buffer on ice for transportation to the lab. The Locke’s buffer contained (mM): NaCl, 145; KCl, 5.4; Na_2_HPO4, 2.2; NaH_2_PO4, 0.9; glucose, 5.6; HEPES, 10 (pH 7.3, adjusted with NaOH). The glands were perfused with Locke’s buffer, then infused with Locke’s buffer containing collagenase P (1.5 mg/ml, Roche), trypsin inhibitor (0.325 mg/ml, Sigma) and bovine serum albumin (5 mg/ml, Sigma), and incubated at 37 **°**C for 20 min. The digested medulla was minced in Locke’s buffer, and filtered through a 100 nm nylon mesh. The filtrate was centrifuged (48 x *g*, 5 min), re-suspended in Locke’s buffer and re-centrifuged until the supernatant was clear. The final cell pellet was re-suspended in pre-warmed DMEM medium (Gibco) supplemented with 10% fetal bovine serum (Gibco) and plated onto poly-L-lysine (0.005 % w/v, Sigma) and laminin (4 µg/ml, Sigma) coated glass coverslips.

Cells were transfected by electroporation using Basic Primary Neurons Nucleofector Kit (Lonza), according to the manufacturer’s protocol and plated onto glass coverslips with mouse Laminin coating over PDL layer (Neuvitro). The cells were incubated at 37 °C with 9 % CO_2_ and used within 48 hours.

### Fluorescent dyes and plasmids for chromaffin cells

For FFN511 (Abcam) imaging, cells were bathed with FFN511 (5-10 μM) in 37°C incubator for 20 min and images were performed after washing out FFN511 in the bath solution. Atto 655 (A655, Sigma) was included in the bath solution at the concentration of 30 μM. PH-EGFP (phospholipase C delta PH domain attached with EGFP) was obtained from Dr. Tamas Balla. PH-mNeonGreen (PH_G_) was created by replacing the EGFP tag of PH-EGFP with mNeonGreen (Allele Biotechnology)^47^.

For knockdown of endogenous NSF in bovine chromaffin cells, a siRNA duplex for bovine NSF (5′-CCAGAUUGUCGAUGUGUUU-3′) labeled with cyanine 3 (si-NSF) and scrambled control siRNA (si-Ctrl) labeled with cyanine 3 were purchased from Sigma-Aldrich. For rescue experiments, si-NSF and a plasmid containing wildtype NSF and mCherry (for recognition of the transfected cell) (Addgene #84334) were transfected into chromaffin cells. The cDNA encoding NSF_E329Q_ (Genecopoeia) was subcloned into EBFP2-C1 (Addgene #54665), where EBFP2 was used to recognize transfected cells.

### Western blot

Total protein was extracted from cultured chromaffin cells or hippocampal cultures using RIPA buffer containing protease inhibitor cocktail (Millipore Sigma). Equal amounts of proteins, determined by BCA protein assay (Invitrogen) were loaded onto 4%–12% Bis-Tris gel (Invitrogen). Proteins were transferred onto PVDF membrane and immunoblotted with the indicated primary antibodies at 4 °C overnight. Membranes were incubated with HPR-labeled secondary antibodies at 22-24 °C for 2 hours and visualized using Bio-Rad ChemiDoc Imaging System. Primary antibodies included anti-NSF (1:2000, Abcam), mouse anti-CHC (1:500, Abcam), rabbit anti-dynamin (1:1000, Cell Signaling Technology), mouse anti-AP2 (1:1000, ThermoFisher Scientific), and β-actin (1:3000; Abcam).

### Electrophysiological recording at chromaffin cells

The method has been described before^22,28,47^. At room temperature (20 ̶ 22°C), whole-cell voltage-clamp and capacitance recordings were performed with an EPC-10 amplifier together with the software lock-in amplifier (PULSE 8.74, HEKA, Lambrecht, Germany). The holding potential was -80 mV. For capacitance measurements, the frequency of the sinusoidal stimulus was 1000 Hz with a peak-to-peak voltage ≤ 50 mV. The bath solution contained 125 mM NaCl, 10 mM glucose, 10 mM HEPES, 5 mM CaCl_2_, 1 mM MgCl_2_, 4.5 mM KCl, 0.001 mM TTX and 20 mM TEA, pH 7.3 adjusted with NaOH. The pipette (2 – 4 MΩ) solution contained 130 mM Cs-glutamate, 0.5 mM Cs-EGTA, 12 mM NaCl, 30 mM HEPES, 1 mM MgCl_2_, 2 mM ATP, and 0.5 mM GTP, pH 7.2 adjusted with CsOH. These solutions pharmacologically isolated calcium currents. For stimulation, we used a 1-s depolarization from the holding potential of -80 mV to +10 mV (depol_1s_). We used this stimulus because it induces robust exo-endocytosis as reflected in capacitance recordings. Since prolonged whole-cell recording slows down endocytosis, we limited to 1 depol_1s_ per cell.

### Confocal imaging at chromaffin cells

Imaging of PH_G_, FFN511, and A655 was performed with an inverted confocal microscope (TCS SP5II, Leica, Germany, 100× oil objective, numerical aperture: 1.4)^24,47^. PH_G_ was excited by a tunable white light laser at 515 nm (laser power set at ∼1-4 mW); FFN511 was excited by an Argon laser at 458 nm (laser power set at ∼2-4 mW); A655 was excited by an HeNe laser at 633 nm (laser power set at ∼12-15 mW); their fluorescence was collected at 520-600 nm, 465-510 nm, and 650-800 nm, respectively. Confocal imaging area was ∼70 - 160 μm^2^ at the XY plane with a fixed Z-axis focal plane∼100-200 nm above the cell-bottom membrane (XY/Z_fix_ scanning). Images were collected every 40-80 ms at 40-60 nm per pixel.

### Fusion modes, close-fusion and pre-close detection with confocal microscopy

Full-fusion was identified as the sudden appearance of PH_G_ spot or ring together with the sudden appearance of an A655 spot, due to PH_G_ and A655 diffusion from the plasma membrane (PM) and the bath into the fusion-generated Ω-profile (fs-Ω, Figure 4) at cell-bottom. FFN511 (pre-loaded in vesicles) fluorescence (F_FFN_) decrease concurrently at the same spot as PH_G_ fluorescence (F_PH_) and A655 fluorescence (F_655_) increased, while measurements was made for estimation of FFN511 release rate. The fusion pore closes at ∼0.05 - 30 s later (close-fusion, Fig. 4), maintains an open pore (stay-fusion, Fig. 4), or shrinks to merge with the plasma membrane (shrink-fusion, Fig. 4)^23,24,29,47^.

Close-fusion was detected as F_655_ (strongly excited) dimming due to pore closure that prevented bath fluorescent A655 from exchanging with bleached A655 in vesicle, while F_PH_ (weakly excited) sustained or decayed with a delay that reflected vesicle pinch off (Figure 4); stay-fusion was detected as sustained F_655_ and F_PH_ (Fig. 4); shrink-fusion was detected as parallel increases and decreases of F_655_ and F_PH_ (Fig. 4).

Pre-close was detected with spot F_655_ bleaching with constant F_PH_, due to fusion pore closure of pre-Ω by strong excitation. It is not due to a narrow pore smaller than A655 molecule size, because after spot dimming, bath application of an acid solution cannot quench the pH-sensitive VAMP2-EGFP or VAMP2-pHluorin overexpressed at the same spot, indicating that the spot is impermeable to H^+^ or OH^-^, the smallest molecules, and thus is closed^22^.

### Imaging data selection of chromaffin cells

For chromaffin cell recorded with a Cm and ICa recording, the data collected within the first 2 min after the break-in of the whole-cell configuration were used, which avoided rundown of endocytosis and Ca^2+^ influx (gradual disappearance of endocytosis and ICa) as previously reported under the whole-cell configuration for a long time^28,48^. For avoiding large fluctuations from individual cells in data analysis, chromaffin cells with less than 5 fusion events were not used. Electrophysiological data (Cm and ICa) were analyzed with Igor Pro (WaveMetrics). Confocal images were analyzed with LAS X (Leica) and ImageJ. Fluorescence intensity from an area covering the fluorescence spot was measured at every image frame.

### Statistical tests

Data were expressed as mean ± s.e.m. Replicates are indicated in results and figure legends. n represents the number of cells, fusion events, or experiments as indicated in the results and figure legends. The statistical test used is the t-test. Although the statistics were performed based on the number of cells, fusion events, and pre-close, each dataset was replicated from at least four primary chromaffin cell cultures. Each culture was from at least three glands from two bovines.

