## supplemental file for "NSF is required for diverse endocytic modes by promoting fusion and fission pore closure in secretory cells"

**Supplementary Information**

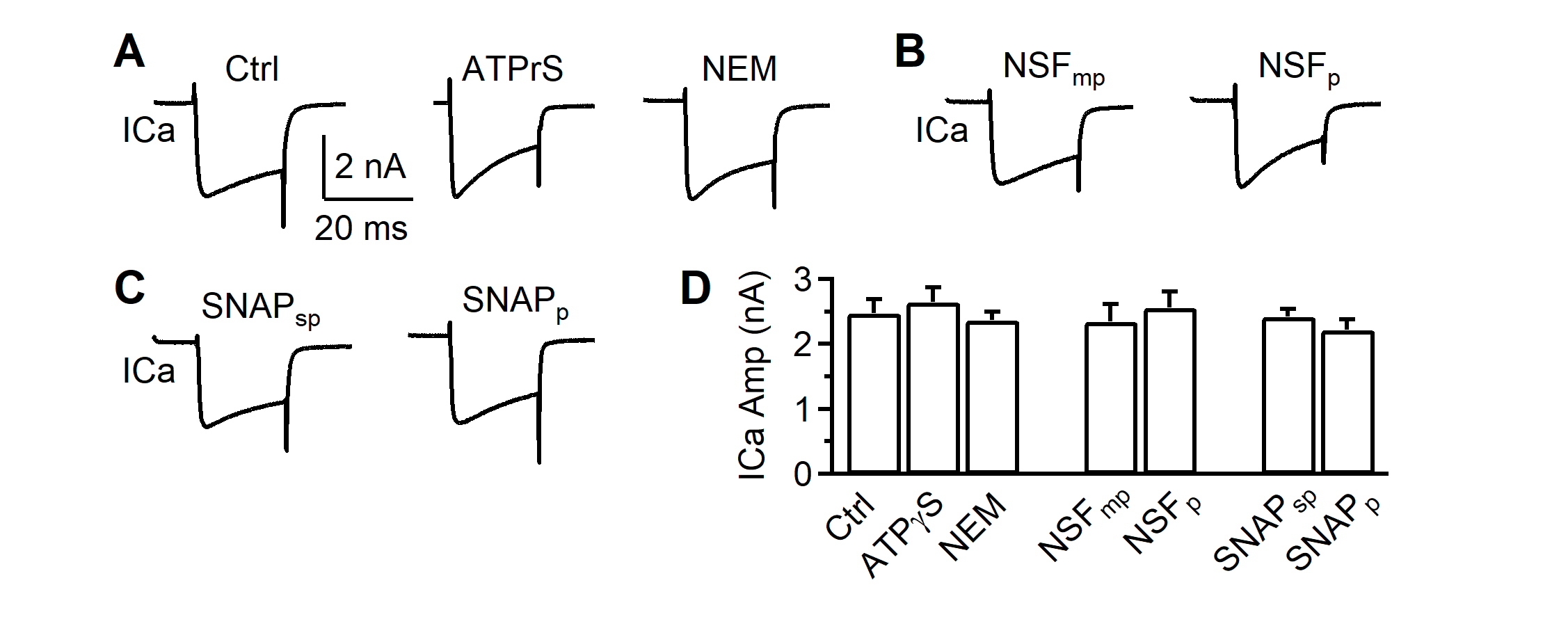

**Figure S1.** NSF blockers do not affect calcium currents and do not cause significant asynchronous release after stimulation; related to Figure 1.

(A)	Sampled traces (single trace) of similar calcium currents (ICa) induced by a 20 ms depolarization from -80 to +10 mV with a pipette solution containing a control solution, 4 mM ATPγS (replacing ATP) or 1 mM NEM. Data were taken in 4 – 10 min after whole-cell break in. Scale bars apply to A-C.

(B)	Similar to panel A (similar ICa), but with NSF_mp_ (1 mM) or NSF_p_ (1 mM).

(C)	Similar to panel B (similar ICa), but with SNAP_sp_ (1 mM) or SNAP_p_ (1 mM).

(D)	Similar amplitudes of ICa induced by depol_20ms_ at 4 - 10 min after break in with a pipette containing the control solution (Ctrl, n = 11), ATPγS (4 mM, n = 6), NEM (1 mM, n = 12), NSF_mp_ (1 mM, n = 10), NSF_p_ (1 mM, n = 9), SNAP_sp_ (1 mM, n = 7) or SNAP_p_ (1 mM, n = 8).

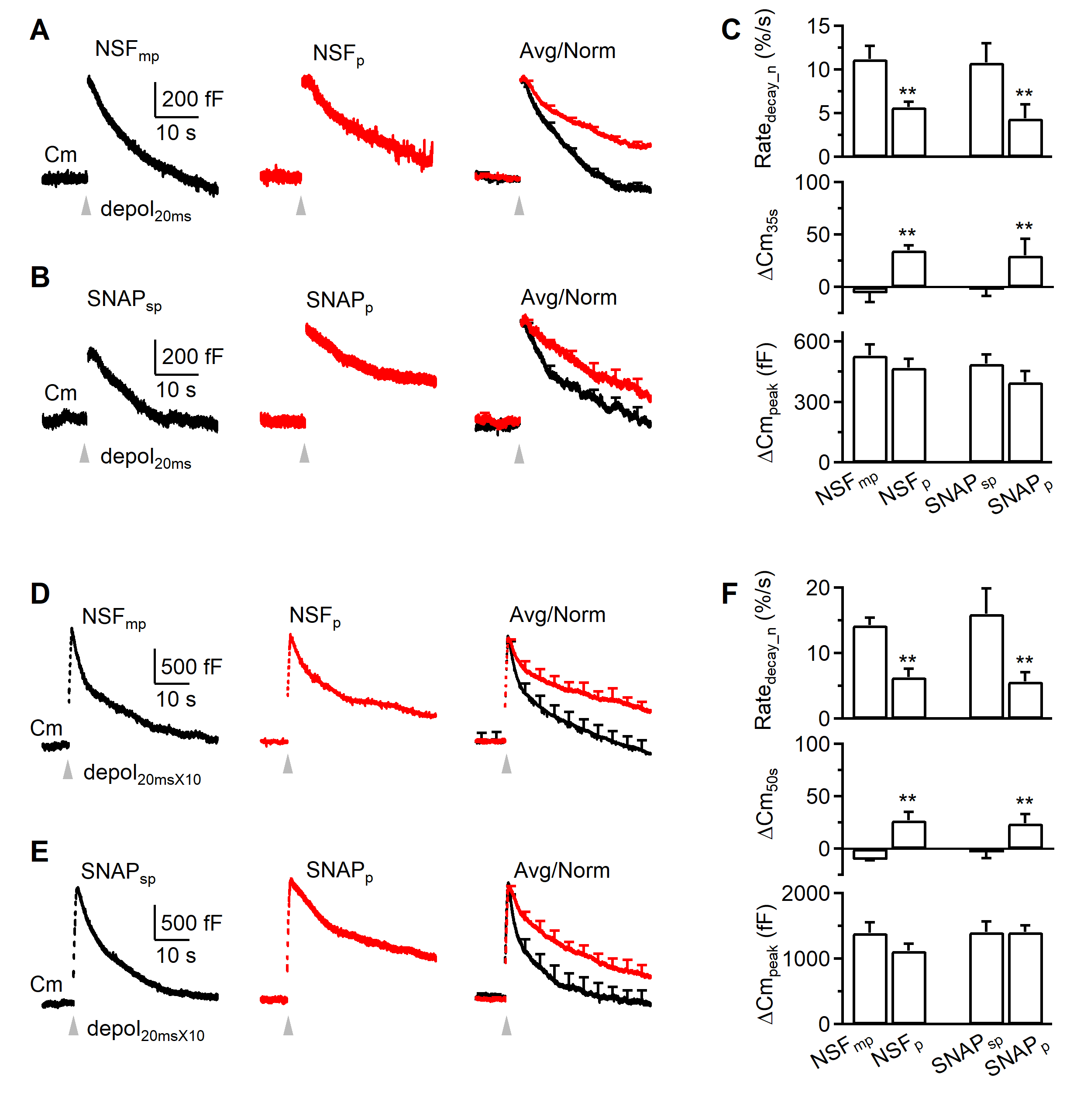

**Figure S2.** NSF is involved in slow and rapid endocytosis at calyces: results observed 2-4 min after whole-cell break-in

(A)	Sampled (single traces, left two traces) or averaged (right, avg) capacitance changes (Cm) induced by depol_20ms_ (arrow) at 2 - 4 min after break in with a pipette containing NSF_mp_ (1 mM, black, n = 10) or NSF_p_ (1 mM, red, n = 9). Traces on the left and middle are single traces, whereas traces on the right are averaged traces with the amplitude normalized for comparison of the decay.

(B)	Similar to panel B, but with SNAP_sp_ (1 mM, black, n = 7) or SNAP_p_ (1 mM, red, n = 8).

(C)	The Rate_decay_n_, ΔCm_35s_, and ΔCm_peak_ induced by depol_20ms_ at 2 - 4 min after break in with a pipette containing NSF_mp_ (1 mM, n = 10), NSF_p_ (1 mM, n = 9), SNAP_sp_ (1 mM, n = 7) or SNAP_p_ (1 mM, n = 8).

(D-F)	Similar arrangement as panels A-C (including the calyx number), respectively, except that the stimulus was depol_20msX10_ and the ΔCm_35s_ was replaced with ΔCm_50s_ (E).

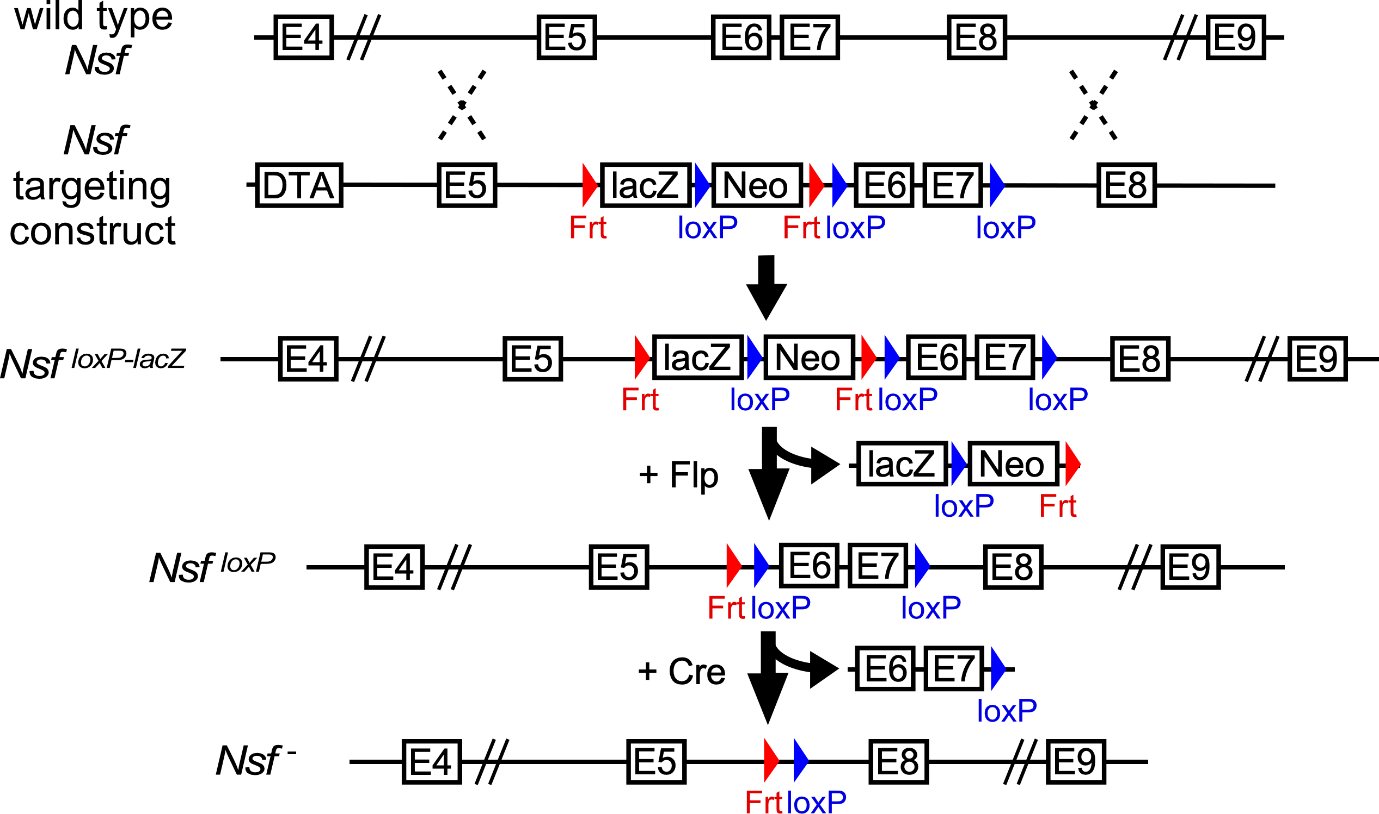

**Figure S3.** Generation of *NSF* conditional knockout allele. In the *NSF* targeting construct created by The Knockout Mouse Project (KOMP), a 5.8 kb *NSF* fragment containing exon 5 and a 3.8 kb fragment containing exon 8 were used as 5’ and 3’ homologous arms, respectively. A synthetic 7.1 kb fragment containing Frt-flanked lacZ reporter gene, Neo cassette, and loxP-flanked Exon 6 and 7 was inserted between the 5’ and 3’ homologous arms. The targeted *NSF^loxP-lacZ^* embryonic stem cells (ESCs) in C57BL/6N background were generated and were purchased from The KOMP repository and were injected into C57BL/6J blastocysts to generate chimeric founder mice. The founder mice were bred with wild type C57BL/6J to generate *NSF^loxP-lacZ^* heterozygotes and Rosa26-Flpe (Jackson stock#009086) mice were used to remove the lacZ and Neo sequences to generate *NSF^loxP^* mice. Crossing *NSF^loxP^* with Cre mice results in *NSF^-^* mice by deleting loxP-flanked exon 6 and 7 and resulting in a shift in the reading frame and premature termination codon in exon 8.

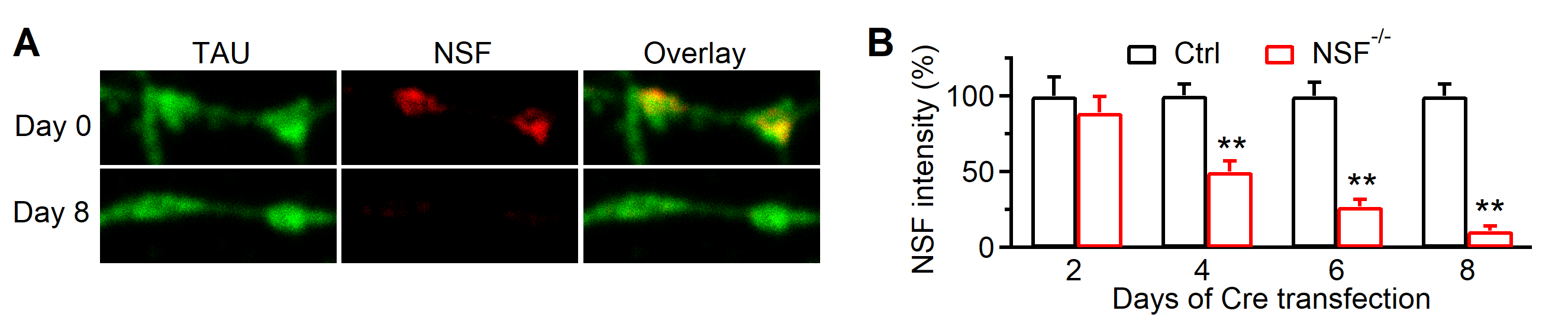

**Figure S4.** Immunostaining showing deletion of NSF in the NSF^-/-^ hippocampal culture

(A) Immnuostaining of TAU (labelling the axon) and NSF in day 0 and day 8 after Cre transfection to *NSF^LoxP/LoxP^* hippocampal cultures.
(B) NSF immunostaining intensity in day 2, day 4, day 6, and day 8 after Cre-mCherry transfection to *NSF^LoxP/LoxP^* hippocampal cultures (red, NSF^-/-^, 3 cultures) or wildtype hippocampal cultures (black, Ctrl, 3 cultures). **: p < 0.01, two-way ANOVA.

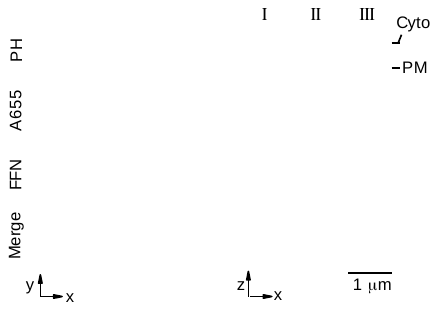

Figure S5. Confocal images of PH_G_, A655, and FFN511 (near cell-bottom) showing pre-spot I, II and III at the XY- (left, ring-shape) and XZ-plane (right, Ω-shape). Data taken from Ref. (1) with permission.
